# Genome wide screen of RNAi molecules against SARS-CoV-2 creates a broadly potent prophylaxis

**DOI:** 10.1101/2022.04.12.488010

**Authors:** Ohad Yogev, Omer Weissbrod, Giorgia Battistoni, Dario Bressan, Adi Naamti, Ilaria Falciatori, Ahmet C. Berkyurek, Roni Rasnic, Myra Hosmillo, Shaul Ilan, Iris Grossman, Lauren McCormick, Christopher C. Honeycutt, Timothy Johnston, Matthew Gagne, Daniel C. Douek, Ian Goodfellow, Gregory J. Hannon, Yaniv Erlich

## Abstract

Expanding the arsenal of prophylactic approaches against SARS-CoV-2 is of utmost importance, specifically those strategies that are resistant to antigenic drift in Spike. Here, we conducted a screen with over 16,000 RNAi triggers against the SARS-CoV-2 genome using a massively parallel assay to identify hyper-potent siRNAs. We selected 10 candidates for *in vitro* validation and found five siRNAs that exhibited hyper-potent activity with IC50<20pM and strong neutralisation in live virus experiments. We further enhanced the activity by combinatorial pairing of the siRNA candidates to develop siRNA cocktails and found that these cocktails are active against multiple types of variants of concern (VOC). We examined over 2,000 possible mutations to the siRNA target sites using saturation mutagenesis and identified broad protection against future variants. Finally, we demonstrated that intranasal administration of the siRNA cocktail effectively attenuates clinical signs and viral measures of disease in the Syrian hamster model. Our results pave the way to development of an additional layer of antiviral prophylaxis that is orthogonal to vaccines and monoclonal antibodies.

## Introduction

Covid-19 has been one of the world’s worst pandemics in modern times. While vaccines have been a major triumph, there is an urgent need to expand the arsenal of preventative measures to address some of their shortcomings^1^. First, virtually all licensed vaccines target the Spike protein^2,3^, converging on a single point of failure exposed to escape mutants and emerging virulent variants^4–8^. Moreover, as all monoclonal antibody (mAb) treatments target this same protein, such antigenic shifts not only hamper the protection of vaccines, but can also reduce the efficacy of a wide range of other treatments. Second, multiple studies have shown that vaccines’ protection, including against severe disease, typically wanes within just a few months, after the second^9^, third^10,11^, or the fourth dose^12^. Third, recent lines of evidence in mice and NHP suggest that updated versions of vaccines have diminished efficacy and may be subject to original antigenic sin^13,14^. These data suggest the limited utility of vaccine updates for emerging VoCs. Finally, several studies consistently show that it is challenging to achieve high protection in immunocompromised individuals, even after repeated dosing^15^, implying that the individuals who most need the vaccine are the ones least likely to benefit from it. Finally, infections in immunocompromised individuals can be prolonged^16,17^, which increases the risk of hyper-evolution and the emergence of VoCs, resulting in major risks to public health.

Backed by the success of multiple previous studies where small interfering RNAs (siRNAs) were effectively used as antivirals^18–21^, we envision that intranasally (i.n.) administered siRNAs are particularly well suited as a vaccine augmentation measure for infections of the upper respiratory tract, where they can mitigate transmission. To this end, we screened over 16,000 RNA interference (RNAi) triggers targeting the SARS-CoV-2 genome in order to identify hyper-potent candidates. The screen relied on a massively parallel assay, Sens.AI, that uses a synthetic biology system to recapitulate the silencing activity of each siRNA candidate against the virus. In our previous studies^22,23^, we used an earlier version of Sens.AI to identify hyper-potent siRNAs against HIV and HCV. However, the previous design took over 6 months of operation. In the new design, we used a quicker method that enhances the signal-to-noise ratio by employing statistical learning in lieu of laborious experimental steps. Extensive computational analyses and *in vitro* experiments yielded a cocktail of two hyper-potent siRNA candidates, effective against all tested viral strains. Intranasal administration of this siRNA cocktail was confirmed as effective in an *in vivo* experiment in the Syrian hamster model of SARS-CoV-2.

## Results

### Screening for hyper-potent shRNA against SARS-CoV-2

We parsed the SARS-CoV-2 genome into a series of potential short hairpin RNA (shRNA) targets (**Supplemental Figure 1**). This process was conducted by tiling the genome with overlapping 50 nucleotide-long sequences, each shifted by a single nucleotide from the other. The target region for the shRNA is a stretch of 22 nucleotides located in the middle of the 50 nucleotide sequence, and the rest of the flanking sequence serves to preserve the genomic context. We then applied multiple *in silico* filters to exclude target regions with low synthesis fidelity, do not pass a minimal threshold of conservation across viral strains, have sequence attributes that are typically associated with poor shRNA response, and those whose seed region can potentially match a human transcripts (**Supplemental Table 1**). In total, this process retrieved 16,471 shRNAs candidates targeting the SARS-CoV-2 genome and sgRNA1 negative strand. In addition, we added to this library a set of 1,118 positive and negative control shRNAs that had been identified in previous screens against cancer-related genes in the mouse genome^22^.

We synthesised these 17,589 shRNAs and their corresponding 50 nucleotide target regions using a DNA oligo pool by Twist Bioscience. Each of these oligos was 185 nucleotide long, and consisted of two PCR annealing sites, the miR-30-based shRNA, a guide and its passenger strand based on our design, a spacer with cloning sites, and a 50 nucleotide region that recapitulates the target site with its genomic context (**Figure 1A**). We used a series of cloning steps to introduce a Venus reporter gene to the spacer region, such that the 3’UTR of Venus included the 50 nucleotide target region, and inserted this entire construct into our retro-vector library (**Figure 1B**).

**Figure 1:**
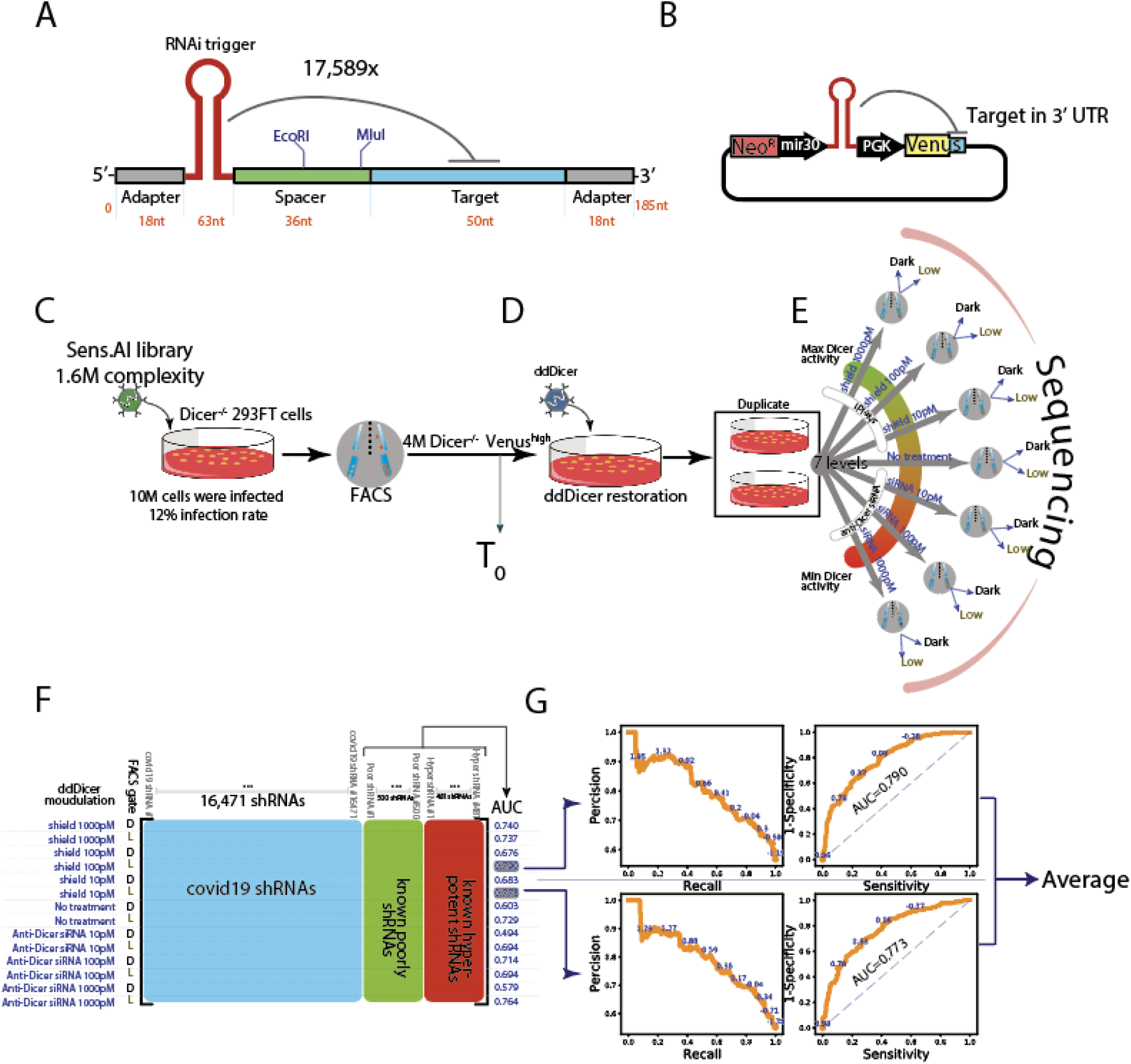
Genome wide screen of SARS-CoV-2 by Sens. **AI (A) Oligo design.** Each oligo had a unique RNAi trigger in the context of a miR-30 backbone along with a 50 nucleotide stretch from the viral genome that includes the RNAi target site **(B) Library design.** Each plasmid contained a specific RNAi trigger and its matching target-site in cis as part of the 3’UTR of a Venus reporter **(C-E) Scheme of a screen in human cells (C) The RNAi machinery is turned off.** Dicer^-/-^ 293FT cells were infected by the plasmid library and sorted for Venus^high^ expression, which formed the T_0_ read out **(D) The RNAi machinery is turned on.** We restored the RNAi machinery by ectopic expression of a destabilised Dicer (ddDicer) **(E) Modulating the activity of the RNAi machinery.** Two biological replicates were subjected to seven different conditions, designed to titer Dicer expression either down, by using anti-Dicer siRNA, or up, by using Shield-1. Upon FACS sorting, Venus^low^ and Venus^dark^ cells were collected and their oligo constructs were sequenced **(F) The resulting data matrix of the screen**. Each row vector represents a specific combination of a treatment condition (the far left column) and a FACS gate (denoted as D, for Dark, and L, for Low). These row vectors describe the enrichment of each tested RNAi trigger (blue) compared to negative (green highlighted) and positive (red highlighted) controls. The AUC column was calculated best on the ability to separate the positive from the negative controls **(G) Selecting the two best conditions.** The two row vectors with the highest AUC were selected. Precision-recall (left) and receiver operating characteristics (right) curves for the intrinsic controls are shown. The screen score of each RNAi trigger is the average of these two row vectors.

Our screening procedure consisted of two steps to reduce the effect of position variation that may be introduced by retro-vector integration. We first started the screen in a human *Dicer*^null/null^293FT cell line, engineered by CAS9/CRISPR knockout (**Figure 1C; Supplemental Figure 2**). The absence of *Dicer* prevents the maturation of shRNAs, effectively uncoupling between the Venus expression and the potency of each encoded shRNA. Overall, we transduced 1.2 million *Dicer*^null/null^ 293FT cells with retro-vectors that encoded our library at a multiplicity of infection (MOI) of 0.8. Three days post-infection, we FACS-sorted four million Venus^high^ *Dicer*^null/null^ 293FT cells out of 50 million cells. These cells represent instances of successful construct integration into genomic loci, resulting in adequate Venus expression. We then restored the expression of *Dicer* by using a synthetic construct and modulated DICER expression to couple between the optical signal and the potency of each shRNA (**Figure 1D**). The synthetic construct was a fusion between human *Dicer* and a destabilising domain (*ddDicer*) that was based on a mutant human *FKBP12* protein, enabling us to dial up the activity of Dicer by using Shield-1^24^. In addition, we employed an siRNA against Dicer to induce the opposite effect, reducing its expression. The principal idea behind these various conditions was to identify a regimen in which the RNAi machinery allows hyper-potent shRNAs to inhibit their targets, but cannot support the activity of less potent shRNAs.

In total, we screened the library across eight different conditions of *ddDicer* expression (**Figure 1E**). The first condition, which we assigned as T_0_, was devoid of *ddDicer* and reflected the non-manipulated relative abundance of the various shRNAs. The other conditions contained increased-doses of either Shield-1 (to induce dd*Dicer*-activation) or anti-Dicer siRNA (to inhibit Dicer). Each of these seven conditions was conducted in two biological replicates. We sorted cells in each replicate into three bins based on their Venus expression: high, low, and dark, followed by sequencing of the low and dark bins in order to decrypt the level and identity of the shRNAs. We also sequenced T_0_ unsorted shRNAs to depict the distribution of the shRNAs in the initial library.Overall, we obtained (2_[replicates]_ x 2_[sorting bins]_ x 4_[Dicer Expression levels]_ + 1_[T_0_]_ =) 17 sequencing libraries with Illumina MiSeq, each composed of 150 bp paired-end reads. In total, we obtained 36 million reads on average for each library. We parsed the 50 base-pair region that corresponds to the target from these libraries, annotated them back to their shRNA, and counted the number of unique appearances of each shRNA in each condition. Finally, we used DESeq225 to measure the enrichment of the shRNA in each condition versus T_0_ and averaged the two biological replicates. This process yielded an 8-by-17,589 matrix (**Figure 1F**), where each column represents an shRNA, for which each row represents one of the treatments (siRNA or Shield-1, each from one of two sorting bins, low and dark), and the enrichment statistic (the right-most column) is represented by AUC as calculated by DESeq2.

Next, we used our internal controls to identify the optimal parameters that separate hyper-potent shRNAs from the rest of the library (**Figure 1F**). We calculated the area under the curve (AUC) for separating the poorly potent from the hyper-potent controls using the DESeq2 enrichment statistic. In general, this process showed that the low Venus bins substantially outperformed the dark bins in terms of separating hyper-potent from poor ones. In addition, the Shield1 containing conditions performed better than the Shield-null conditions, with 10 pM and 100pM concentrations being the best performers. Therefore, we decided to focus on these two conditions with the low Venus gate as the optimal ones to separate the hyper-potent shRNA molecules from the poorly performing ones. After focusing on RNAi triggers with high sequencing coverage, these two conditions had AUC scores approaching 80% for the internal controls. More importantly, these two conditions displayed perfect positive predictive value for the internal controls when restricting the recall to the top 5% of the list.

After identifying the optimal parameters, we ranked the candidate SARS-CoV-2 shRNAs using a process similar to the one employed for the internal controls. For each shRNA, we used the DESeq2 statistic for each of the conditions under the same sequencing coverage restrictions. The end result was a rank-ordered of shRNAs across all tested conditions, with the most potent shRNA at the top, to the least potent one at the bottom.

### Validation of screen results

Next, we validated the ranking of our screen using multiple methods. First, we analysed the correlation between the SARS-CoV-2 shRNA statistic tests across the two top performing conditions. This analysis found a Pearson correlation between the reported statistics of the two conditions to be 72.2% (p<10^-9^), showing that the screen has significant internal consistency (**Figure 2A**). Next, we focused on the sequence features of our shRNAs. Previous studies reported that highly potent shRNAs are typically associated with the absence of adenine in the 20th position of the guide^22^. Therefore, we evaluated the frequency of adenine as a function of the averaged screen score from the two conditions. This analysis revealed a significant correlation (Pearson=90.4%, p<10^-20^) between the frequency of SARS-CoV-2 shRNAs without adenine in their 20th position and the average screen statistic (**Figure 2B**). In fact, the top shRNAs in our screen were virtually all depleted of adenine in position 20. Finally, we compared the results of our screen to *in-silico* shRNA potency predictions by a published machine learning algorithm^26^ (**Figure 2C**). While the prediction of these algorithms is far from being perfect for each individual RNA trigger, we found a highly significant correlation (Pearson=90.6%, p<10^-20^) between its scores and the average screen score.

**Figure 2:**
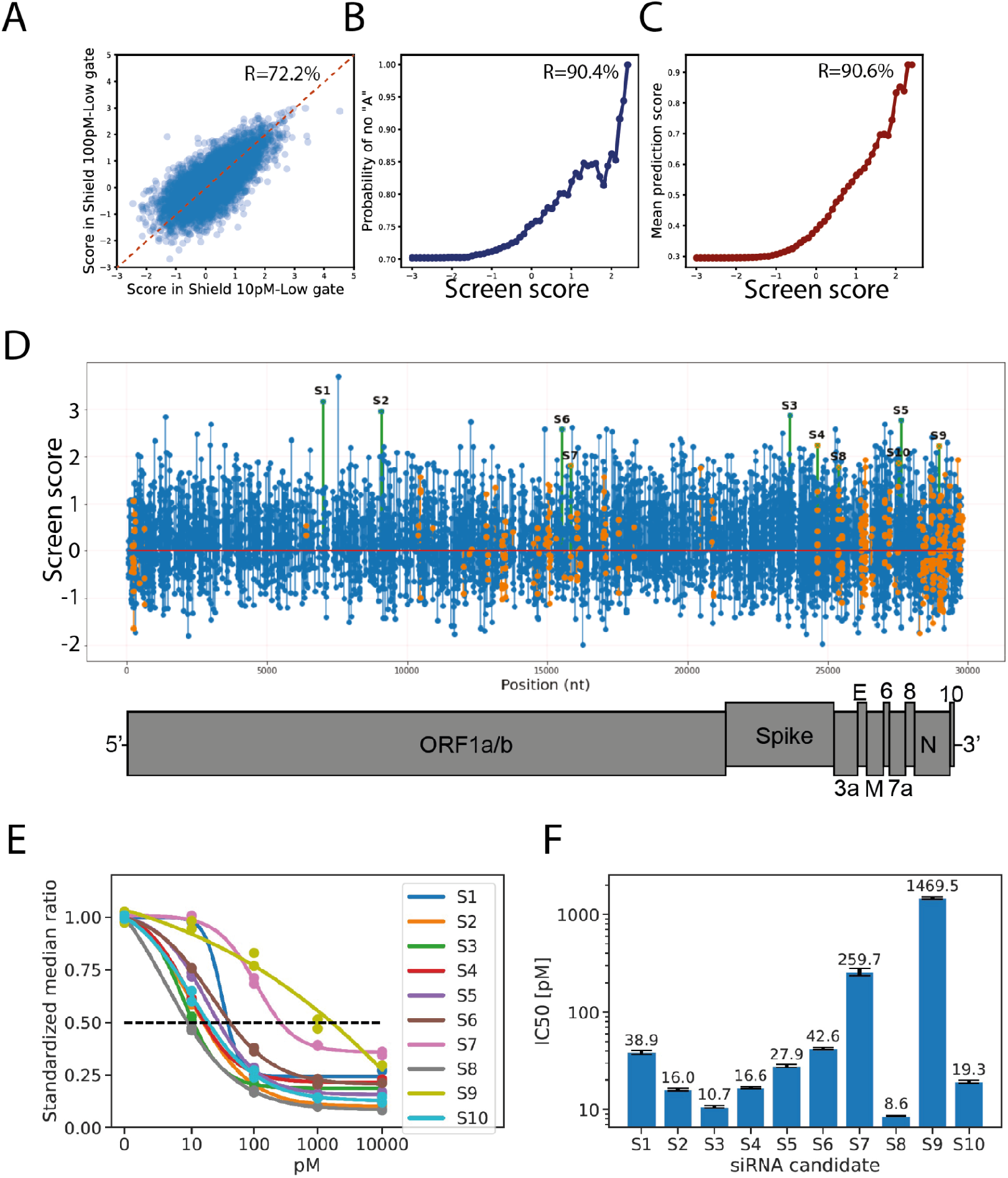
Validation of screen results. **(A) The screen has high internal consistency.** The results show a Pearson correlation of 72.2% between the screen scores in the two top performing conditions **(B) Enrichment of features associated with potent shRNAs.** The probability of *not* finding an Adenine in position 20 of the RNAi triggers targeting SARS-CoV-2 increases with screen scores, recapitulating previous studies indicating that when adenine occupies position 20 it weakens shRNA maturation **(C) The screen’s resultant scores are highly correlated with bioinformatic predictions.** The screen scores, representing enrichment statistics as calculated by DESeq2, are highly correlated with a published machine learning algorithm predicting the potency of shRNAs **(D) Screen results and the selected candidates**. The graph shows the resultant screen scores of each RNAi trigger that passed our quality threshold, as a function of position along the viral genome. Orange: RNAi triggers that target conserved regions between SARS-CoV and SARS-COV-2. Green: The selected 10 candidates. Blue: all other RNAi riggers **(E) The dose-response curves of the selected 10 candidates.** siRNAs were tested using a reporter assay. The plot represents the standardised median ratio between the expression of mCherry (reporter gene) and GFP (control gene) **(F) Potency scores of each of the 10 selected candidates.** The IC50 value of each of the 10 candidates was calculated based on the dose-response curves in (E).

Encouraged by these results, we manually selected ten candidates for further experimentation (**Table 1; Figure 2D**). The first five candidates (S1-S4 and S6) were selected mainly based on their average screen score in the two best performing conditions while trying to span multiple virus genes. The other five candidates (S5 and S7-S10) were also selected based on their screen scores, but restricted to 22mer regions that are fully conserved between SARS and SARS-CoV-2, as we hypothesised that these regions might be applicable tofuture spillovers of beta-coronaviruses as well. To test our shortlisted shRNAs, we cloned their target region into the 3’UTR of mCherry and converted each shRNA to its corresponding siRNA. We then transfected each reporter with decreased doses of its corresponding siRNA to identify its half maximal inhibitory concentration (IC50). Eight of the ten siRNAs tested exhibited IC50 values below 50pM (**Figure 2E-F; Supplemental Table 2**), five of which demonstrating IC50 values below 20pM. Only one candidate, S9, showed a relatively poor IC50 value in this assay (IC50 >1.4uM). Overall, these results show that our novel genome-wide screen method identifies hyper-potent siRNAs within a single cycle.

**Table 1:** The best performing top ten shRNAs identified by the Sens.AI screen that were selected for further validations and development. ^+^Indicates whether the target site is also conserved in SARS-CoV, in addition to SARS-CoV-2.

| Label | Strand | Gene | Screen score | SARS-CoV<sup>+</sup> | IC50 [pM] |
| --- | --- | --- | --- | --- | --- |
| S1 | + | nsp3 | 3.18 | No | 38.9 |
| S2 | + | nsp4 | 2.97 | No | 16.0 |
| S3 | + | Spike | 2.87 | No | 10.7 |
| S4 | + | Spike | 2.24 | Yes | 16.6 |
| S5 | + | ORF7A | 2.77 | No | 27.9 |
| S6 | + | RdRP | 2.58 | No | 42.6 |
| S7 | + | RdRP | 1.82 | Yes | 259.7 |
| S8 | - | Spike | 1.79 | Yes | 8.6 |
| S9 | + | N | 2.23 | Yes | 1469.5 |
| S10 | - | ORF7A | 1.87 | Yes | 19.3 |

### Discovery of siRNAs against SARS-CoV-2 using a bioinformatic pipeline

In parallel to the Sens.AI screen, we employed a more traditional discovery pipeline to identify siRNAs against SARS-CoV-2. The main reason for this pipeline was to assess the performance, technical and logistical properties of our novel Sene.AI in comparison to the state-of-the-art siRNA prediction algorithms. To this end, we used three open-source siRNA potency-predicting algorithms: RNAxs^27^, DSIR^28^, OligoWalk^29^, to computationally evaluate over 815 potential target sites, focusing on regions that previously showed good results against SARS-CoV ^30–32^. There was a relatively low correlation between the algorithms, with Spearman correlations between 0.4 and 0.02 (**Supplementary Figure 3**) and we manually selected siRNA candidates that were consistently better in all programs, excluding candidates with seed regions complementary to the human transcriptome. This list yielded 88 siRNAs that we synthesised and tested by the same reporter assay described above. First, we subjected 1nM of each candidate to a reporter assay. This analysis revealed that the average repression was ~45% (s.e. 0.26) (data not shown). We then prioritised the most promising 27 siRNAs and retested them at 500pM and 100pM concentrations (**Supplementary Figure 4**), finding that 9 of these 27 candidates inhibit the reporter expression by more than 50% at 100pM.

### siRNAs conferring protection against multiple SARS-CoV-2 variants

Next, we assessed our top candidates from the sensor screen and the bioinformatic pipeline using a gold standard live SARS-CoV-2 *in vitro* infection assays. We transfected Vero E6 cells with 100nM of each of the Sens.AI siRNA candidates in triplicates. As a negative control, we also transfected cells with a mock siRNA that targets eGFP. After 24 hours, we challenged the cells with either 60xTCID50, 600xTCID50, or 6000xTCID50 of live SARS-CoV-2 (ancestral strain). Finally, we measured the level of viral load 48 hours after the initial infection via qPCR probing the RdRP and the E genes.

We found that six out of the nine tested siRNAs from our screen were able to dramatically lower the amount of viral RNA. While the results were qualitatively consistent across all three virus titers tested (**Figure 3 and Supplementary Figure 5**), we decided to focus on the 600xTCID50 titer for future experimentations, because it yielded the greatest dynamic range (**Figure 3A**). In these conditions, our best performing five siRNAs repressed genomic viral load by over 95%. Interestingly, both S8 and S10 showed weak responses, at the level of ~10% SARS-CoV-2 inhibition compared to the control siRNA, despite very high potency in the reporter assay (IC50<20pM). However, unlike the other siRNA candidates, these two target the virus negative strand, which is an intermediary in the replication process. Therefore, we hypothesised that targeting this intermediary RNA molecule likely does not interfere with viral replication. To further confirm our top candidates, we measured suppression of live virus by another method, using quantitative media (**Figure 3B**). Again, we found a strong inhibitory activity of the top three out of four siRNAs, with viral repression reaching about two orders of magnitude compared to the GFP siRNA control. We next used the same settings to test 5 of the most potent siRNAs from the open-source discovery method. Only one one candidate, Hel14, was able to repress the viral load by more than 95% as the top siRNAs from our Sens.AI screen (~90% inhibition) (**Figure 3C**).

**Figure 3.**
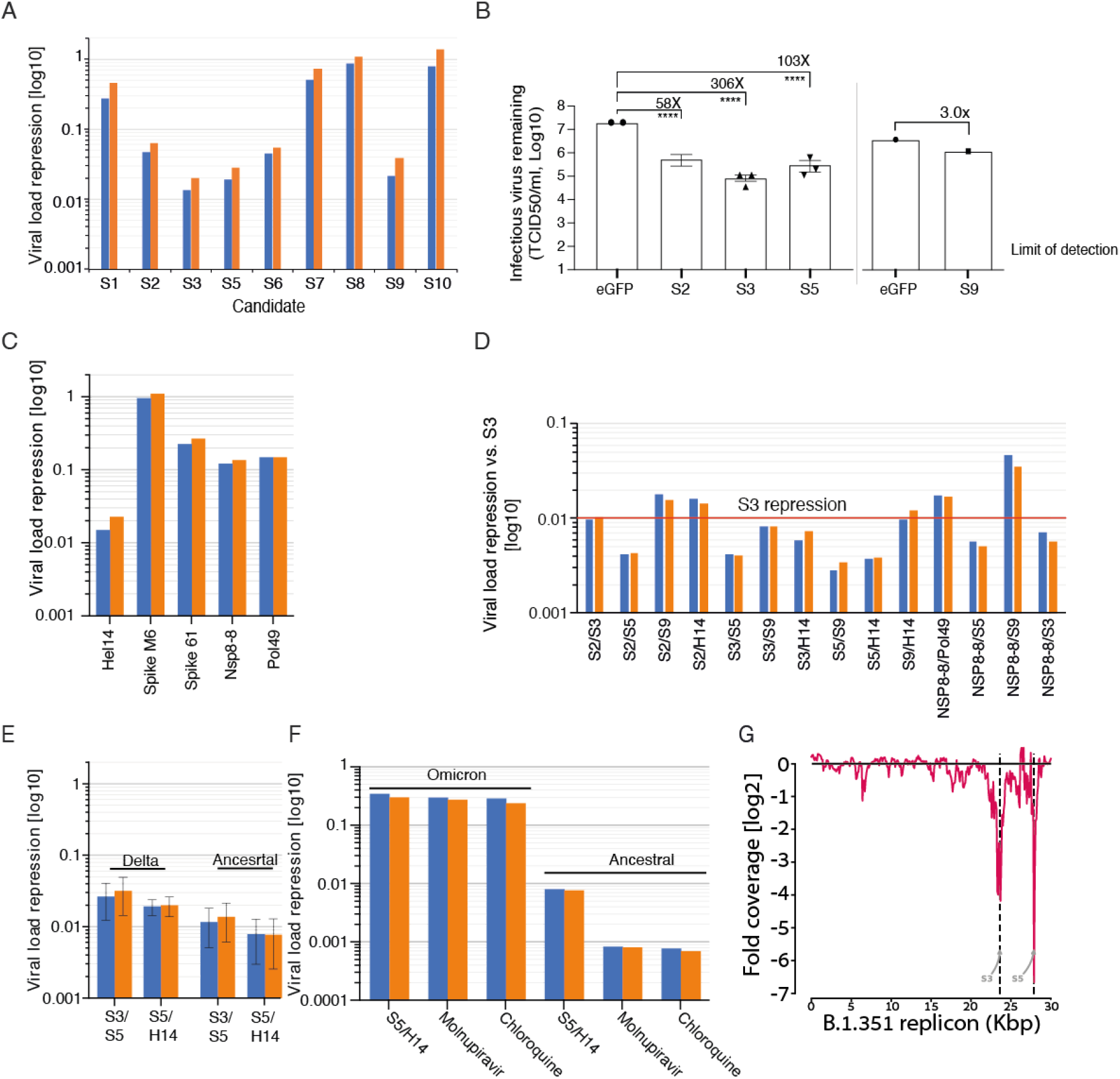
siRNAs repress live SARS-CoV-2 replication in VeroE6 cells. Blue and orange: qPCR results of the E and RdRP transcripts, respectively. Unless noted othewise, the 100% viral load was calibrated to viral level after treatment with an anti-GFP siRNA **(A) The viral load of SARS-CoV-2 (ancestral strain) after treatment with the top siRNA candidates from the Sens.AI screen (B) TCID50 levels of the SARS-CoV-2 (ancestral strain) following treatment with four of the top siRNA molecules.** In each batch of the experiments, an siRNA against eGFP was used as a negative control **(C) The viral load of SARS-CoV-2 (ancestral strain) after treatment with the top siRNA rom the bioinformatic pipeline (D) Testing the effect of various siRNA cocktails against the ancestral strain.** The results were calibrated to the repression of S3 at the same concentration **(E) The viral load against Delta versus the ancestral strain (F) The viral load of Omicron versus the ancestral strain after treatments with S5/Hel14 and other types of anti-virals (G) DeSEQ2 analysis of SARS-CoV-2 replicon treatment with the S3/S5 siRNA cocktail.** We observed a sharp coverage decrease around the S3 (~23.5kbase) and S5 cleavage (~28kbase) sites (FDR values 4×10-83 and 6×10-22, respectively) along the replicon sequence.

Next, we searched for the optimal combination of siRNA pairs out of our best performing candidates. These cocktails included four siRNAs from the Sens.AI screen, augmented by three of the most promising siRNAs from the open-source discovery pipeline. We tested 14 different 2-siRNA cocktails in the live virus assay and compared the results to repression by S3 alone, because as a monotherapy it had shown the highest repression level (**Figure 3D**). We identified five cocktails where the two siRNA components exhibited a synergistic effect, multiplied by several folds in comparison to repression by S3 alone at the same concentration. S5 turned out to be a repeat component in most of these cocktails and we decided to prioritise two of these cocktails moving forward: S5/S3 and S5/Hel14.

Our cocktails are highly resistant to emerging VoCs. First, we tested the cocktails against either the ancestral strain versus the Delta variant. This process showed that the cocktails confer substantial repression, on par with the ancestral strain (over 95% repression) (**Figure 3E**). Interestingly, S5 is tolerant and shows efficacy against the Delta strain, despite the fact that it has a mutation in position 14 of its target site. Second, we tested the S5/Hel14 cocktails against Omicron BA.1 using a similar setting. Similar to other reports^33^, the Omicron variant does not replicate as fast as other VoCs *in vitro*. Since these lower replication rates reduce the dynamic range of our assays, we also added two positive controls, chloroquine or molnupiravir, which are known to be potent inhibitors of Omicron BA.1. The activity of the S5/Hel14 cocktail was indeed diminshed in the Omicron variant. Nevertheless, it inhibited the viral replication to a similar level as induced by the positive control treatments (**Figure 3F**). This suggests that the results are more consistent with lower dynamic range due to slow viral replication rather than reduced potency of the RNAi cocktail. Finally, we tested the activity of the cocktails against our novel replicon system, which recapitulates the function, but not infectivity of the Beta SARS-CoV-2 strain ^34^. Consistently, the cocktails repressed the replicon by 10-15 fold, indicating that they confer a robust inhibition profile across diverse VoCs. Importantly, analysis of high throughput sequencing data showed that cells that were treated with the S3/S5 cocktail had specific depletion of reads at the siRNA cleavage sites (**Figure 3G**). These data provide mechanstic support that the reduction in viral load is directly related to the siRNA silencing.

### High throughput saturation mutagenesis to assess siRNA cross reactivity

Next, we wondered about the cross reactivity of our RNAi strategy against future VoCs. To this end, we developed a saturation mutagenesis assay using our Sens.AI strategy. We repeated the same RNAi-off/RNAi-on strategy as the SARS-CoV-2 screen. The only difference was that in the previous screen that had a variety of shRNA triggers and a perfectly matched target site, whereas in this assay, we fixed the shRNA trigger to be of S5 and created a series of mutated target sites (**Figure 4A**). We used this strategy to screen 2,143 mutations in the S5 target site, exhaustively evaluating virtually every possible two substitution mutations in this site. To the best of our knowledge, this is the largest *in-cellulo* saturation mutagenesis that was tested with an RNAi trigger.

**Figure 4.**
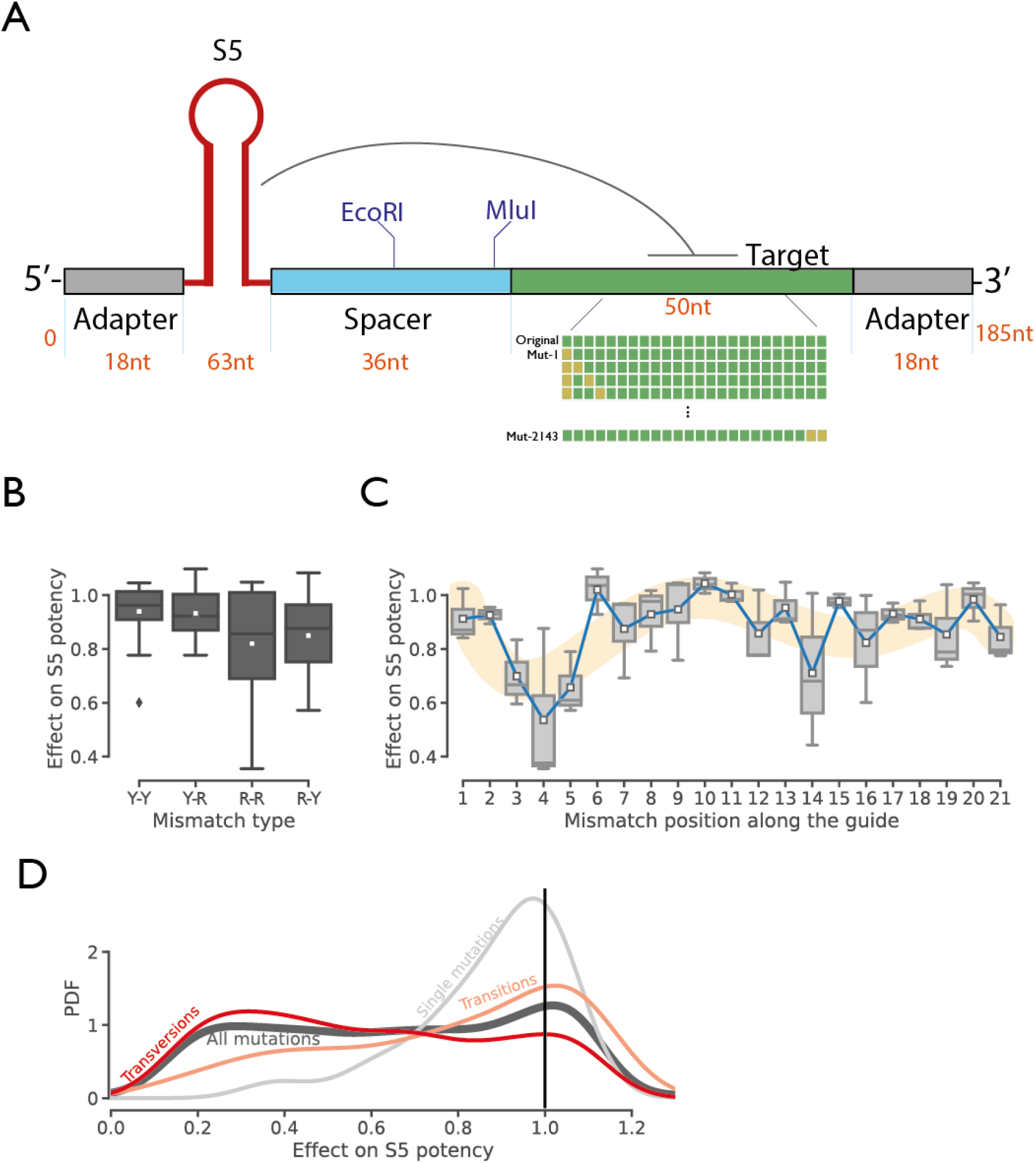
Saturation mutagenesis. **(A) The Sensi.AI oligo design.** The library consisted of the S5 as an shRNA trigger with 2143 mutations, exhausting every possible single- and double-mismatch possibilities in the siRNA target sites **(B) Effect of mutation type on the S5 potency.** The analysis shows all single mutations stratified by the mismatch type (Y-pyrimidine, R-purine; e.g. first letter: guide, second letter: target) **(C) The effect of mismatches by position.** The analysis shows all single mutations stratified by position, where position is based on guide strand orientation. Blue: means, yellow: smoothed mean effect based on position **(D) The distribution of effect of mutations on S5.** The vertical lines signifies no difference than the screen score of a target site without any mutation. Black: the distribution of effects of all 2143 single and double mutations. Grey: the distribution of all single mutations. Orange: the distribution of all double transition mutations. Red: the distribution of all double transversion mutations.

We validated the saturation mutagenesis screen by replicating previous trends about siRNA target mismatches. First, we stratified the results based on the number of mismatches. On average, a single mismatch in the target site reduced the Sens.AI scores by 13% compared to no mismatch. As expected, this figure was significantly smaller (t-test, p<8×10^-9^) than double mismatches, which reduced the Sens.AI scores by 35% on average. Next, we stratified the results based on the type of guide-target mismatch (**Figure 4B**). To avoid confounder effects, we restricted this analysis only to the subset of 63 single mutation instances, which represent every possible substitution in position 1 to 21 of our target site. As described in previous studies^35,36^, purine-purine mismatches had significantly stronger effect (one sided t-test, p<0.02) on diminishing the Sens.AI scores than all other types of mismatches. We also analysed the effect of the position of a single mismatch mutation on the activity of S5 (**Figure 4C**). Similarly, consistent with previous studies^37^, the seed region showed the greatest sensitivity to mismatches, whereas the cleavage site had relatively smaller sensitivity to these mutations.

Overall, our throughput screen shows that the S5 target site can tolerate a wide variety of mutations without a significant loss of potency (**Figure 4D; Supplementary Figure 6**). Most single mutations had up to 8% reduction in the Sens.AI score and about 25% of them even expect to enhance a stronger response, which could be attributed to Ago2 disassociation dynamics^37^. In general, we estimate that S5 would lose most of its potency in the case of a double mutation in the target site. To better understand the dynamics of these double mutations, we compared the estimated effect of double transitions to double transversions.

Our results show that double transitions are much more tolerated. Most double transitions resulted with less than 15% reduction in the Sens.AI score, whereas most double transversions resulted with more than 54% reduction in the Sens.AI score. Based on previous studies, the former mutation type is seven times more common than the latter^38^, suggesting that S5 can tolerate to some extent the prevalent type of double mutations in the target site.

### *In vivo* validation

In order to assess the preventative efficacy of our siRNA cocktails in a disease model of COVID-19, we administered the S5/Hel14 cocktail to Syrian hamsters before a live virus challenge. We decided to use this cocktail over the S3/S5 one since S3 targets the Spike-encoding region, which is prone to mutations as a vaccine- and monoclonal antibody-escape mechanism. We employed a dosing schedule that consisted of one i.n. dose of ~400ug/kg of the siRNA cocktail per day on days −7, −3, and −1 using our propietary lung delivery formulation. In addition, we used the same dosing schedule to treat another group of hamsters with a known and highly potent siRNA against hepatitis

C virus (HCV), as a negative control. As a positive control, we administered bamlanivimab^39^ (LY-COV555), an FDA authorised treatment for COVID-19, intraperitoneally (i.p.) on day −1 to another group of hamsters. Each group had six male hamsters and we challenged them intranasally at day 0 with 4×10^3^ PFUs of the WA-1 ancestral SARS-CoV-2 strain.

Our results show that our siRNA cocktail confers protection against SARS-CoV-2 infection when used as a pre-exposure prophylaxis treatment in the Gold Standard Syrian hamster disease model (**Figure 5A)**. The negative control group experienced an average of >7% weight loss by Day 5 as described in previous studies^40–42^. The siRNA-treated group exhibited significant protection against weight loss compared to the negative control group by day 5 (p < 0.05; bootstrap hypothesis testing and Bonferroni adjusted) (**Figure 5B)**. This weight loss was statistically indistinguishable from the positive control group that was treated with bamlanivimab. In addition, we observed an order of magnitude suppression of the viral load in the lungs of the treatment group by day 5 compared to the negative control based qPCR measurements of the RdRP gene (ΔVL_lung_=10.3x, p_lung_<0.001; bootstrap hypothesis testing and Bonferroni adjusted) (**Figure 5C**). Similarly, we also observed a significant suppression, albeit to a lesser extent, of the viral load in the nares (ΔVL_lung_=2.5x, p_lung_<0.05; bootstrap hypothesis testing and Bonferroni adjusted) (**Figure 5D**). In both cases, the antibody was stronger than the siRNA treatment, suggesting that additional dose adjustments are required for the siRNA treatment.

**Figure 5.**
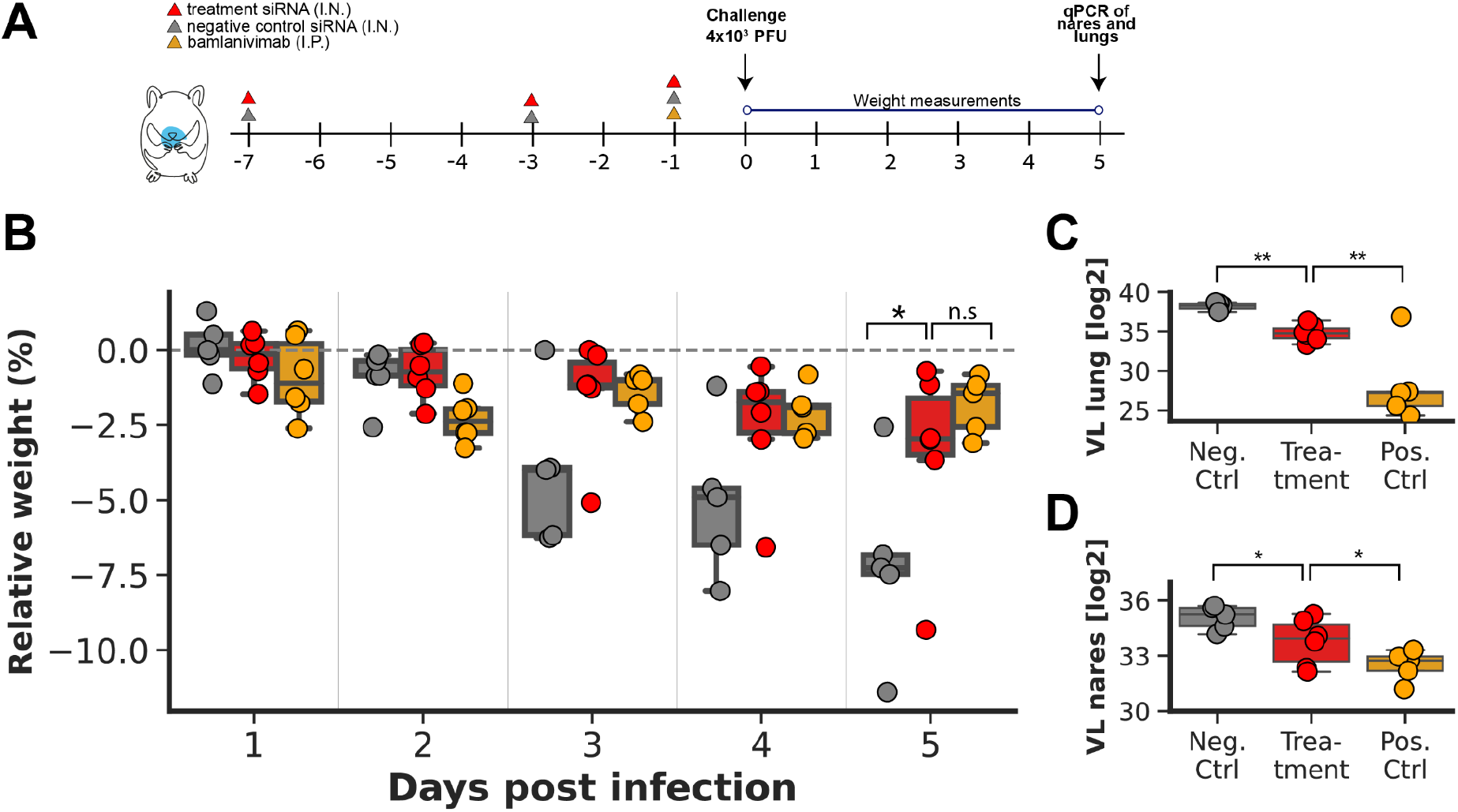
Prophylactic treatment of SARS-CoV-2 infection in Syrian hamsters. **(A) Dosing schedule.** Syrian hamsters pre-treated with a non-targeting siRNA (negative control, grey), the LY-CoV555 antibody (positive control, yellow) or our lead siRNA cocktail (treatment, red) were infected with 4×10^3^ PFUs of the ancestral SARS-CoV-2 virus **(B) Weight change after infection**. The box plot presents the change in weight by treatment group relative to the infection day **(C-D) Viral load (VL) by qPCR.** All measurements are based on qPCR of the RdRP gene five days post infection from either homogenised **(C) lungs (D) nares**. In all panels p value is presented as: *<0.05,**<0.001.

We also tested the same siRNA cocktail with three more treatment variations: with chemically modified siRNAs and i.n. administration (variation #1), with chemically modified siRNAs and by nebulizer administration (variation #2), and no chemical modification and nebulizer administration (variation #3) (**Methods**). Both treatment variation #1 and variation #2 showed significant reduction of the viral load in the lungs (p_lung-variation #1_<0.001; (p_lung-variation #2_<0.001; bootstrap hypothesis testing and Bonferroni adjusted) (**Supplemental Figure 7A**). In addition, we found a smaller but significant reduction of the viral load in the nares (p_lung-variation #1_<0.001; (p_lung-variation #2_<0.05; bootstrap hypothesis testing and Bonferroni adjusted) (**Supplemental Figure 7B**). But none of these variations protected against weight loss by day #5 (**Supplemental Figure 7C**), suggesting that they have slower pharmacokinetics than the primary treatment arm or pose a less tolerable profile by the animals.

Taken together, our results show that siRNA treatment can effectively protect the upper respiratory tract against SARS-CoV-2 infection, significantly attenuating the viral load and passociated clinical manifestations prototypically exemplified by weight loss.

## Discussion

Prophylaxis, particularly the kind resistant to the emergence of novel variants of concern, has been repeatedly identified as a missing component in the arsenal of therapies used to fight the continuous COVID-19 pandemic. In this study, we report, for the first time to our knowledge, on a systematic, genome-wide RNAi screen against SARS-CoV-2. We tested over 16,000 RNAi triggers in a massively parallel reporter assay and validated the best performers in an *in vitro* live virus assay. In addition, we tested 88 siRNAs identified via open-source, *in-silico* discovery methods. We then tested multiple siRNA pairs as cocktail treatments in the live virus assay in order to identify instances where the siRNA components confer synergistic effects when combined. These cocktails proved to be active against three different strains of the virus, namely Beta, Delta and the ancestral one. Finally, we showed the efficacy of these cocktails as pre-exposure preventatives against SARS-CoV-2 infection in Syrian hamsters.

Our study also has certain limitations. First, it is unclear for how long the siRNA cocktails remain effective and thus might be most appropriate in situations where high exposure risk is present, such as the case for front-line workers, immunodepression (transient or chronic), an outbreak in the household, and during the height or an infection “wave”. Second, while our siRNA cocktail protected Syrian hamsters from disease, the effect was mainly observed in the upper respiratory tract. While this is the chief site of infection and shows great promise in terms of mitigating transmission, it will require further adaptation for use in a treatment setting, which would require optimization for delivery to the lower respiratory tract and through to the lung parenchyma. Third, for the purpose of this proof-of-principle study, we adopted a pre-exposure treatment regimen that was composed of multiple days prior to infection. We aim to further optimise the regimen prior to advancing into non-human primates and ultimate clinical trials in humans.

Despite these caveats, the current study substantially contributes to the global efforts to curb COVID-19, as well as future other viral pandemics of similar mortality and morbidity proportions, by illustrating the efficacy of intranasally administered siRNA cocktails as pre-exposure preventatives resilient to the likely mutational evolution of culprit pathogens. This is particularly encouraging given the alarming rate at which we are currently witnessing the emergence of resistance against therapies of the neutralising antibodies and vaccine class in novel SARS-CoV-2 variants. Importantly, our proposed approach is not affected by Spike mutations, nor does it rely on a functional immune system. Instead, it harnesses the natural RNAi pathway, active in the lining epithelial cells of the respiratory tract that are the focal point of infection transmission, and is completely orthogonal to the vaccine and other immuno-modulatory approaches. Finally, the global proportions of the pandemic, and the restrictive cold-chain storage and shipment requirements associated with many of the vaccine and antibody approved therapies, urgently call for a durable solution that would also be accessible to difficult to reach rural, less affluent populations. Our intranasal administration allows for widespread deployment that is not reliant on novel or advanced medical infrastructure for accessibility. We believe that this study illustrates a paradigm shift in the approach to pandemic preparedness, and we expect that our discovery and validation pipeline will be applicable to other emerging pathogenic threats, including novel beta-coronaviruses and the Flu.

## Acknowledgements

This study was funded in part by the Bill & Melinda Gates Foundation and by the intramural program of the National Institutes of Health. The findings and conclusions contained within are those of the authors and do not necessarily reflect positions or policies of the Bill & Melinda Gates Foundation. This work was conducted in part under CRADA 2020-0597 between The VRC/NIAID and Eleven Therapeutics.

## Methods

### Cell cultures

HEK293FT and VeroE6 cells (and their derivatives) were grown in DMEM (Dulbecco’s Modified Eagle Medium), supplemented with 10% Fetal Bovine Serum, 100 U/ml penicillin and 100 μg/ml streptomycin (all Thermo Fisher Scientific), at 37°C with 5% CO2. The HEK293FT cell-line was obtained from Thermo Scientific (cat. No R70007).

### Generation of a *DICER* Knock-Out cell line

The HEK293FT-Dicer Knock-Out (KO) cell line was engineered by CRISPR/Cas9 in the parental 293FT cell-line. The guide RNA (gRNA) had the following sequence: AAGAGCUGUCCUAUCAGAUC. The gRNA was modified with a phosphorothioate modification to prevent nuclease degradation and was obtained from Merck-Millipore. 80 pmol of gRNA were complexed with 4 ug of TrueCut Cas9 Protein (ThermoFisher) and transfected to the cells using the Amaxa nucleofactor. KO efficiency was determined using Sanger sequencing.

### Reporter assay experiments

For each candidate siRNA we extracted its target sequence within a 150 base-pair context. The target sequence was cloned in a plasmid at the 3’UTR of a constitutively expressed mCherry reporter protein. HEK293FT cells were transfected with Lipofectamine 3000 (Life Technologies) according to the manufacturer’s guidelines, either with the sensor alone or with the sensor plus siRNA at the relevant concentration. A second plasmid expressing eGFP was co-transfected at a 1:1 molar ratio, serving as an internal control for transfection efficiency. Cells were collected and analysed 48 hrs post-transfection using a MACSquant VYB flow cytometer (Milteyi Biotec). The potency of each candidate was evaluated as the median ratio of mCherry to eGFP radiance amplitude.

### Cloning of the sensor library

The shRNA-Sensor library was assembled via a two-step procedure. A library of ~20,000 oligonucleotides in which each shRNA was joined to its cognate Sensor by a linker harbouring EcoRI and MluI restriction sites was obtained from Twist Bioscience. The library was first PCR-amplified and cloned using XhoI and MfeI in a recipient retroviral vector. The latter contained the Hygromycin resistant miR30, including its 5’ portion (5’mir30). In the second step, a 3’mir30-PGK-Venus cassette was inserted between the shRNA and its Sensor, integrated via the EcoRI and MluI sites in the linker.

### Live virus experiments

Unless indicated differently, 10,000 VeroE6 cells were transfected with 100nM of siRNA treatment and infected with SARS-Cov-2 24 hours post siRNA treatment. Liverpool, Bristol and Delta strains were used at 60xTCID50 and Omicron at 1200xTCID50. 48 hours post infection, 40ul of the media was collected directly into 160ul of TRIzol™ LS Reagent (ThermoFisher). Viral RNA was extracted using the Direct-zol-96 RNA Kits (ZYMO research) and used as a template for qRTPCR to measure viral copy number. Chloroquine Diphosphate (BioVision) was used at a final concentration of 50uM and Molnupiravir (Focus Biomolecules) was used at a final concentration of 20uM. The sequencing of the Beta replicon was described previously^34^.

### SARS-CoV-2 detection

Viral genome copy number was measured by qPCR using the Charité/Berlin Primer Probe Panel (IDT) and the TaqPath™ 1-Step Multiplex Master Mix (ThemoFisher). We then compared the average Ct values of each condition to the Ct values of eGFP siRNA.

### Adaptation of the screen to investigate siRNA mismatch tolerance

The oligos for the S5 mutagenesis saturation assay were obtained from Twist Bioscience and cloned into the shRNA-Sensor library as described above. Similar to the previous screen, we also incorporated 948 control shRNA from Fellmann et al. ^22^, 511 which are highly potent and 437 of low potency. This screen followed a similar structure to the genome-wide screen with a few modifications: (a) we modulated the RNAi machinery with only three conditions: up regulation of Dicer, down regulation of Dicer, and no modulation. A machine learning pipeline was built to distinguish between the potent and weak control shRNA. The best classifier was chosen using cross-validation, and had an 81.1% precision rate on average (s.e. of 1.14%). The classifier assigns a potency score for each trigger-target pair in the assay. The effect of each mutated target was determined as the classifier score for the mutated trigger-target pair over the score of the perfectly matched trigger-target pair.

### In-vivo experiments

Male Golden Syrian hamsters age 6-8 week were treated with siRNA cocktails at days −7, −3 and −1 pre-infection. Intranasal (IN) administration was performed on hamsters in Study using a proprietary formulation. Each hamster was placed on a sterile surgical pad and lightly stretched out to better place a firm grip on the scruff. The hamster was turned on its back to allow the hamster to breathe and be comfortable. With the neck and chin flat and parallel to the pad, the tip of the pipettor was placed near the left nostril of the hamster at a 45-degree angle, and 5 μL of dosing material was administered to the nostril with a 2–3 sec interval in between for a total of 25 μL/nostril. The hamster was held in this position for 5 seconds or until it regained consciousness, then the administration was repeated for the other nostril for a total of 50 μL/hamster. After the procedure, the hamster was returned to its cage and monitored for 5-10 minutes for any adverse reactions.

For nebulizer administrations siRNA were diluted in PBS+Gelatin 0.5 mg/ml. Aerosol was produced using Vibrating Mash Nebulization (VMN), and nebulization was performed in a Biological Safety Cabinet (BSC).

When indicated, 1mg of mAb555 was administered Intravenously at day −1. At day 0 hamsters were infected with SARS-CoV-2 using intranasal infection of 4×10^3^ PFU virus. At day 5 post infection, hamsters were culled and the trachea and lung were collected for further analysis.

We computed all p-values via bootstrap hypothesis testing, by subtracting from each hamster the mean of its group, resampling hamsters from each group with replacement 20000 times, and computing the fraction of samples in which the difference between the means was greater than the real difference. The presented p-values are after Bonferroni-correction for the four arms.

## Supplementary Material

### Supplementary Figures

**Supplementary Figure 1:**
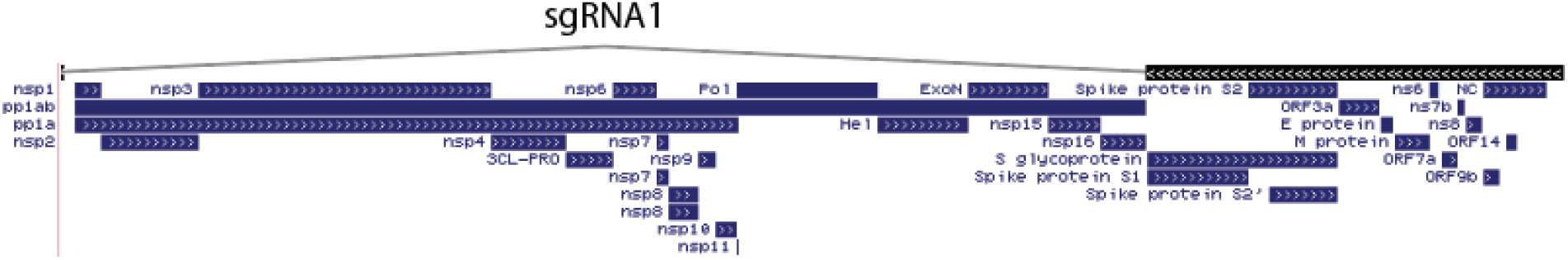
The negative strand sgRNA1 (black) includes all the structural proteins and the leader peptide. Its 5’ position starts from the right most region. In blue: the various proteins in the SARS-CoV-2 genome.

**Supplementary Figure 2:**
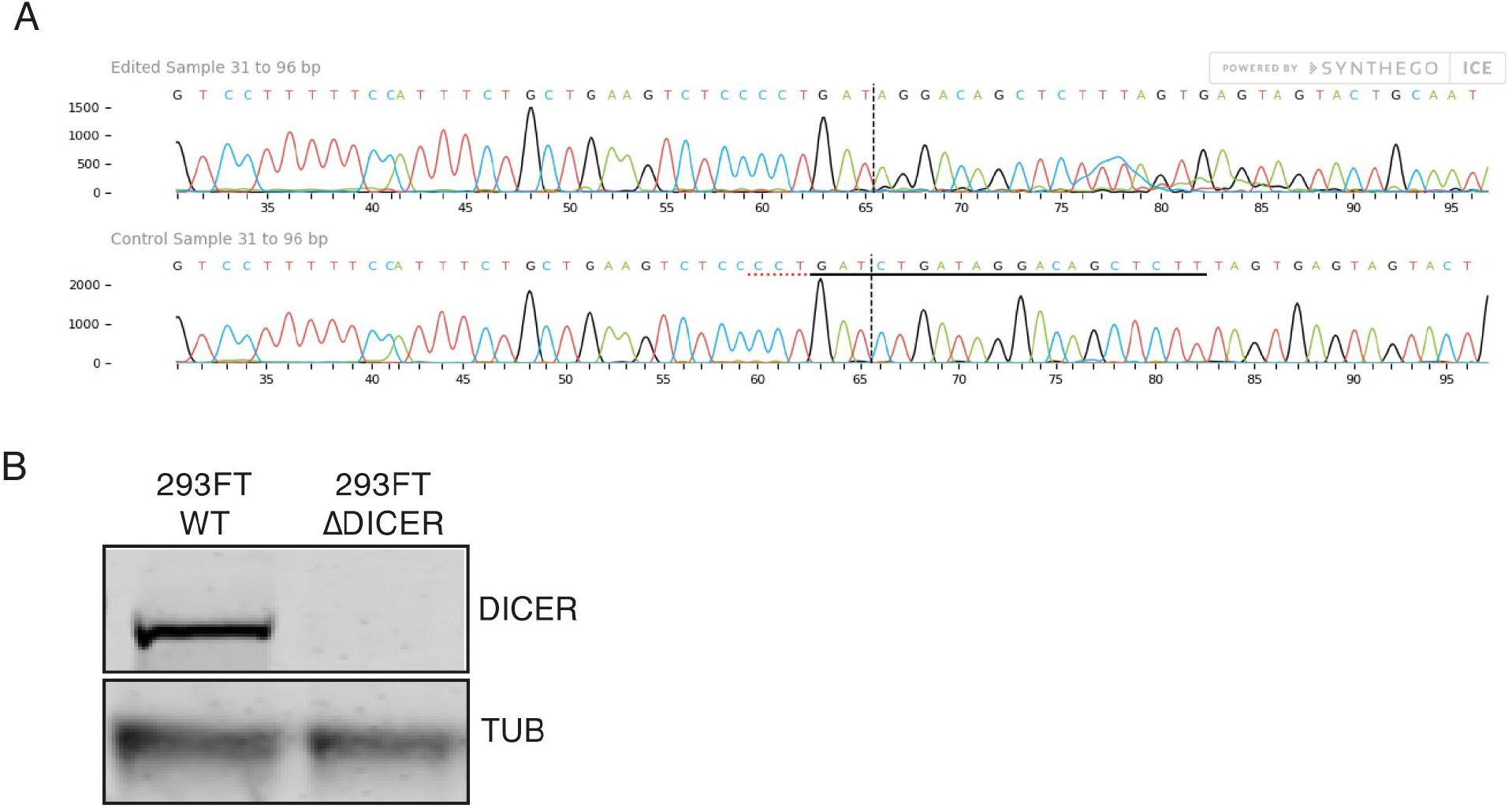
Validation of DICER KO in 293FT cells. **(A)** Sanger sequencing analysis of the gRNA region was performed using the Synthego ICE tool shows a 5 nucleotides deletion. **(B)** Western blot analysis of the parental and KO cells shows complete deletion of DICER protein in the KO cells.

**Supplementary Figure 3:**
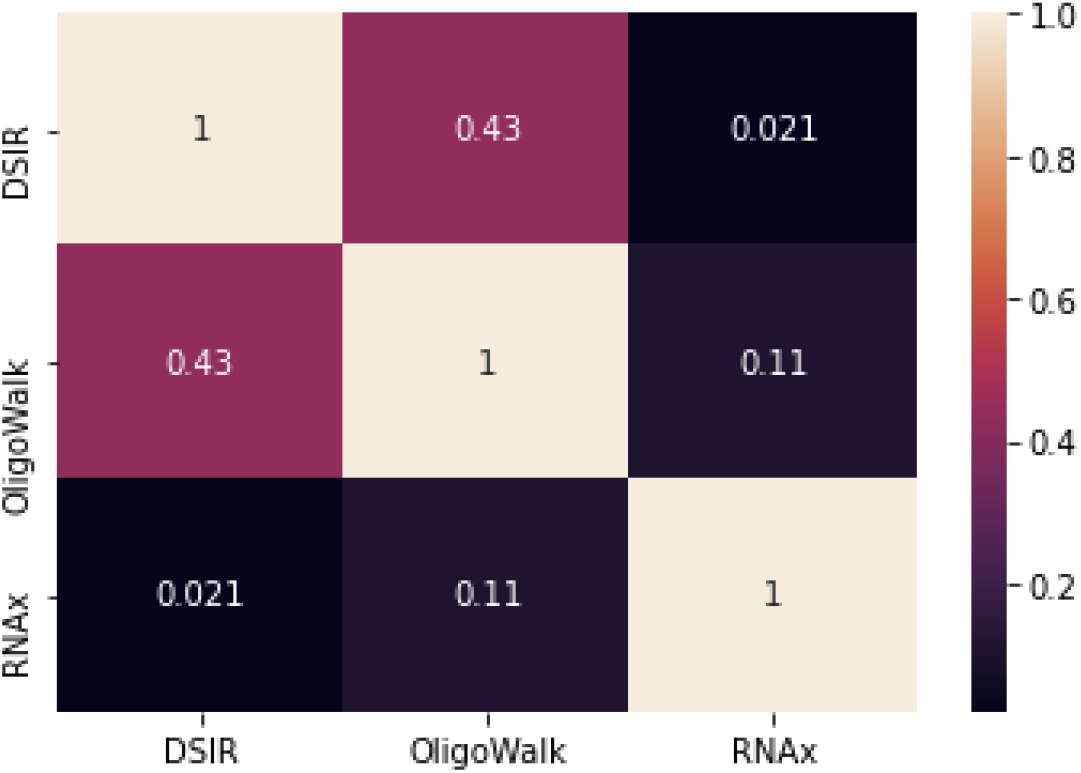
The cross-correlation matrix between DSIR, Oligowalk, and RNAx with respect to the sorted siRNA candidate list.

**Supplementary Figure 4.**
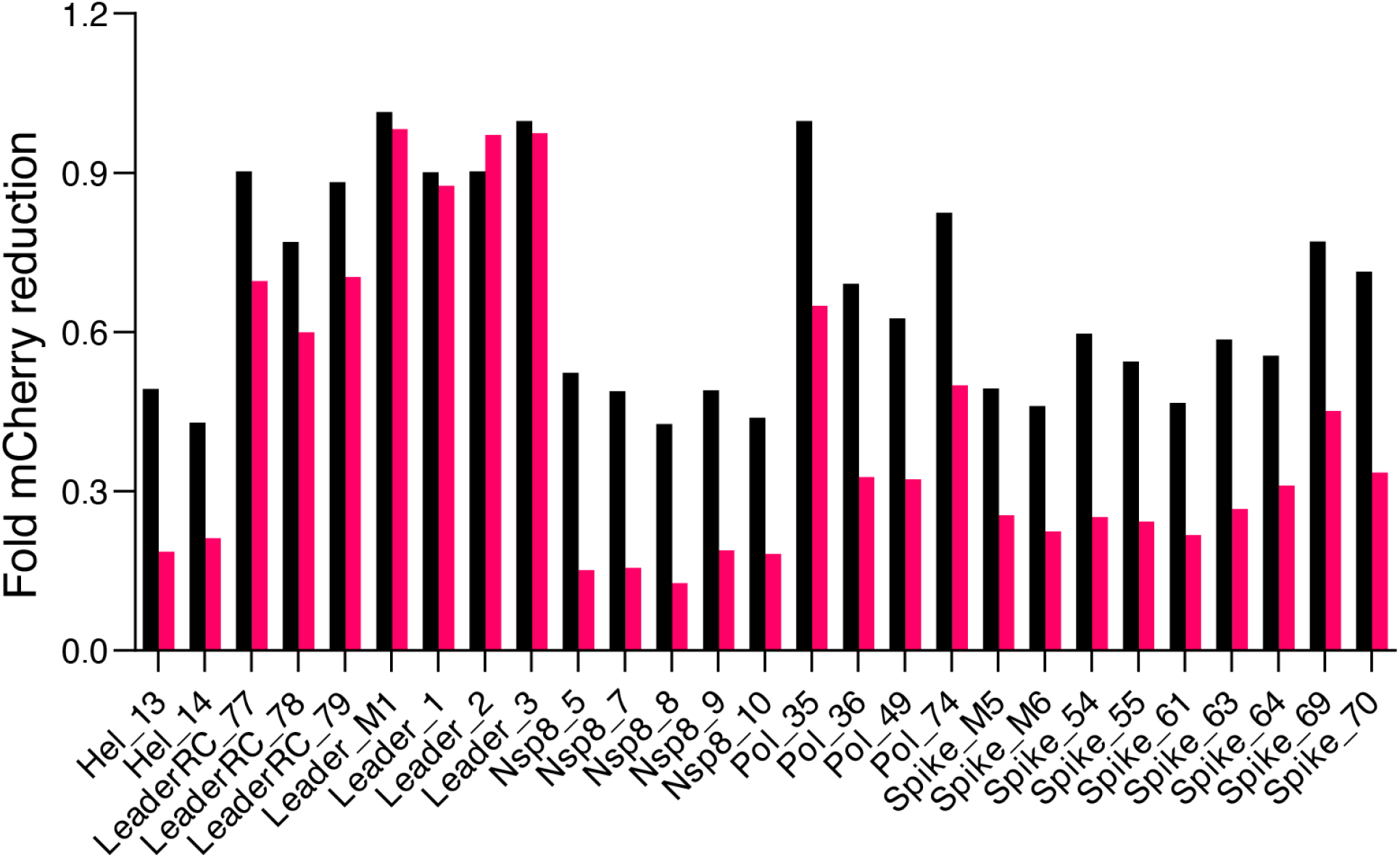
Validation of the selected siRNAs by a reporter assay. Cells were treated with each siRNA at 100pM (black) or 500pM (pink) together with their respective target site fused to the 3’UTR of mCherry. siRNA activity was calculated by measuring the %mCherry positive signal in the GFP positive cell population.

**Supplementary Figure 5:**
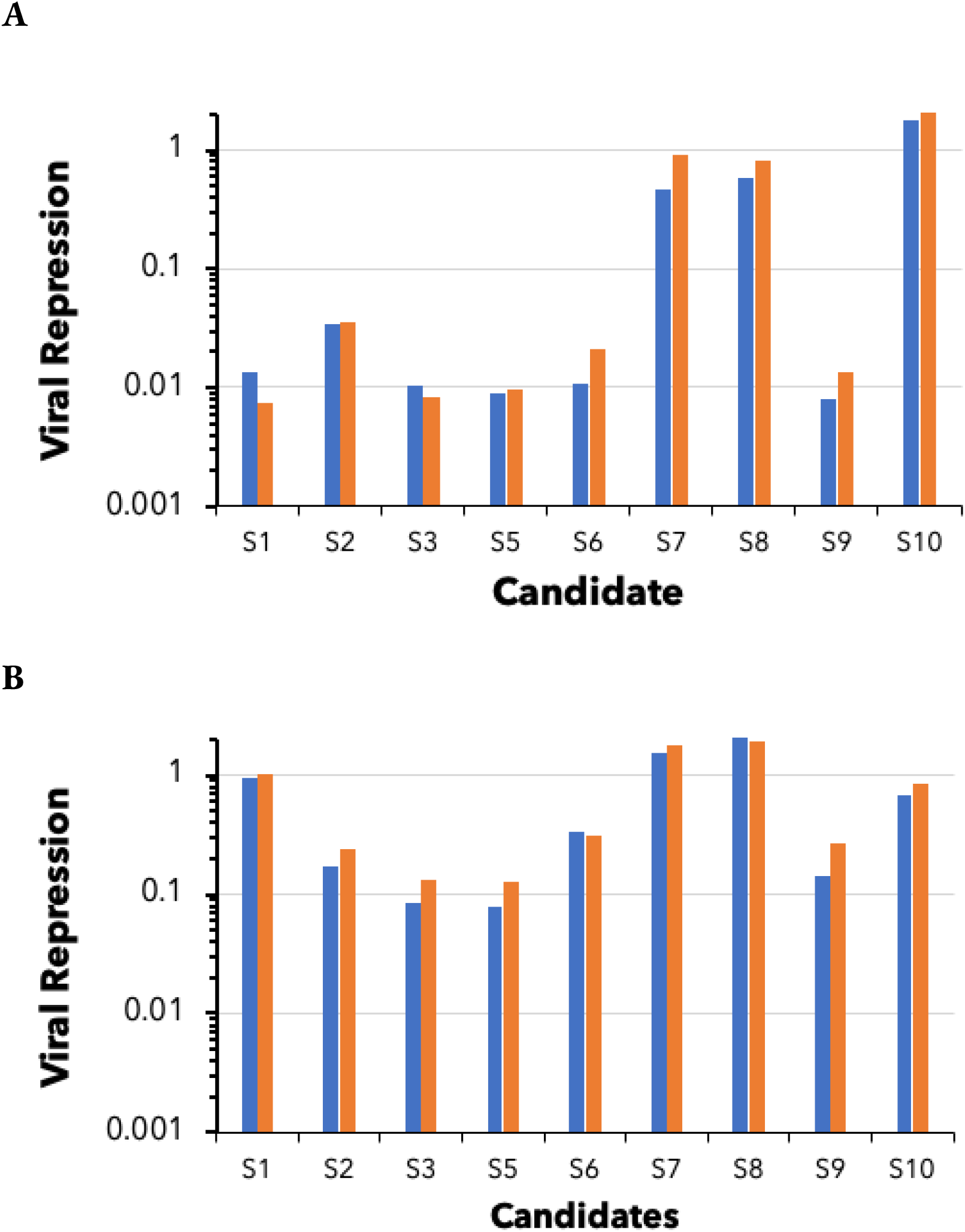
Viral load repressions after an siRNA prophylaxis treatment in VeroE6 cells as measured by qPCR. The level is calibrated to 100% using siRNA against GFP. Blue: RdRP; Orange: E-gene **(A) Challenging the cells with 6000xTCID50 (B) Challenging the cells with 60xTCID50**.

**Supplementary Figure 6.**
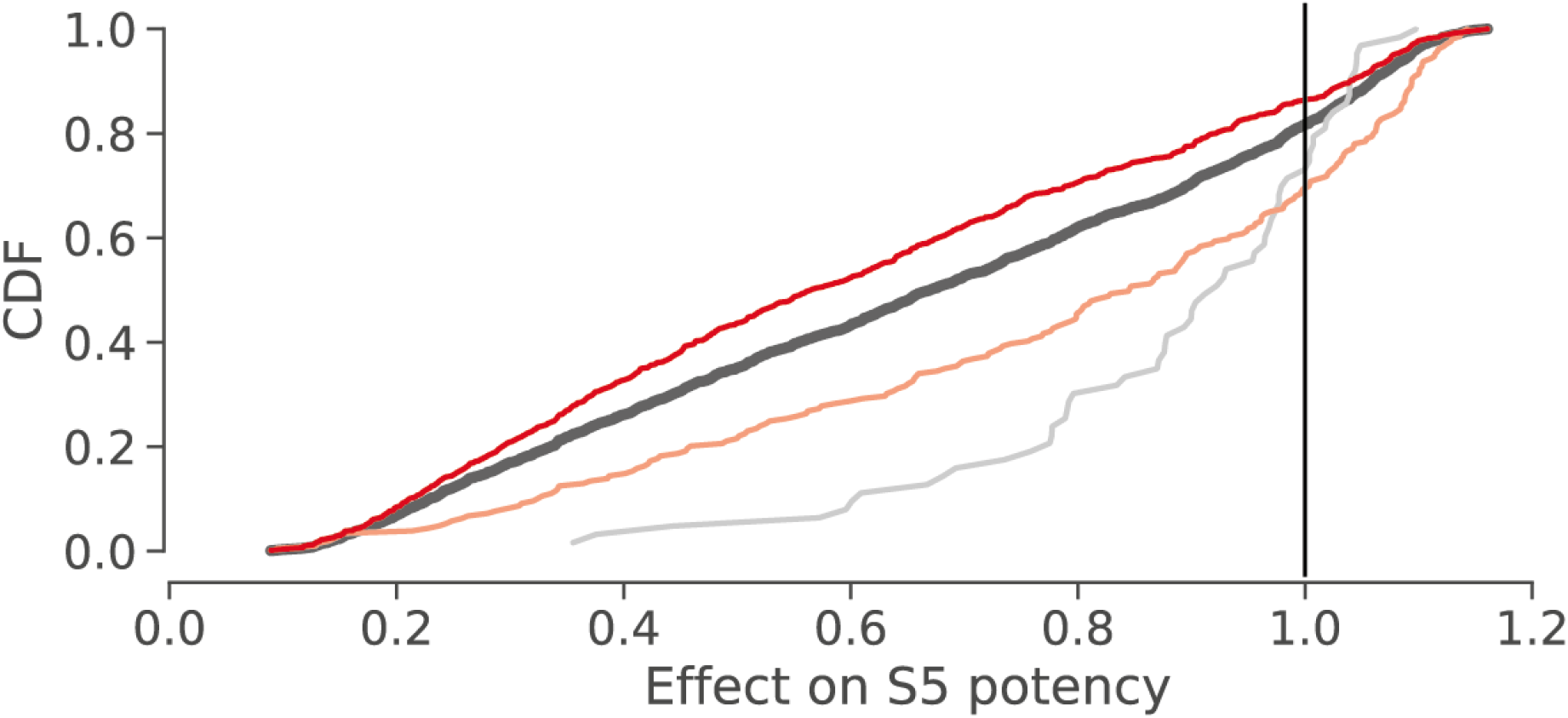
The cumulative distribution function of the effect of mutations on S5. The vertical lines signifies no difference than the screen score of a target site without any mutation. Black: the distribution of effects of all 2143 single and double mutations. Grey: the distribution of all single mutations. Orange: the distribution of all double transition mutations. Red: the distribution of all double transversion mutations.

**Supplementary Figure 7.**
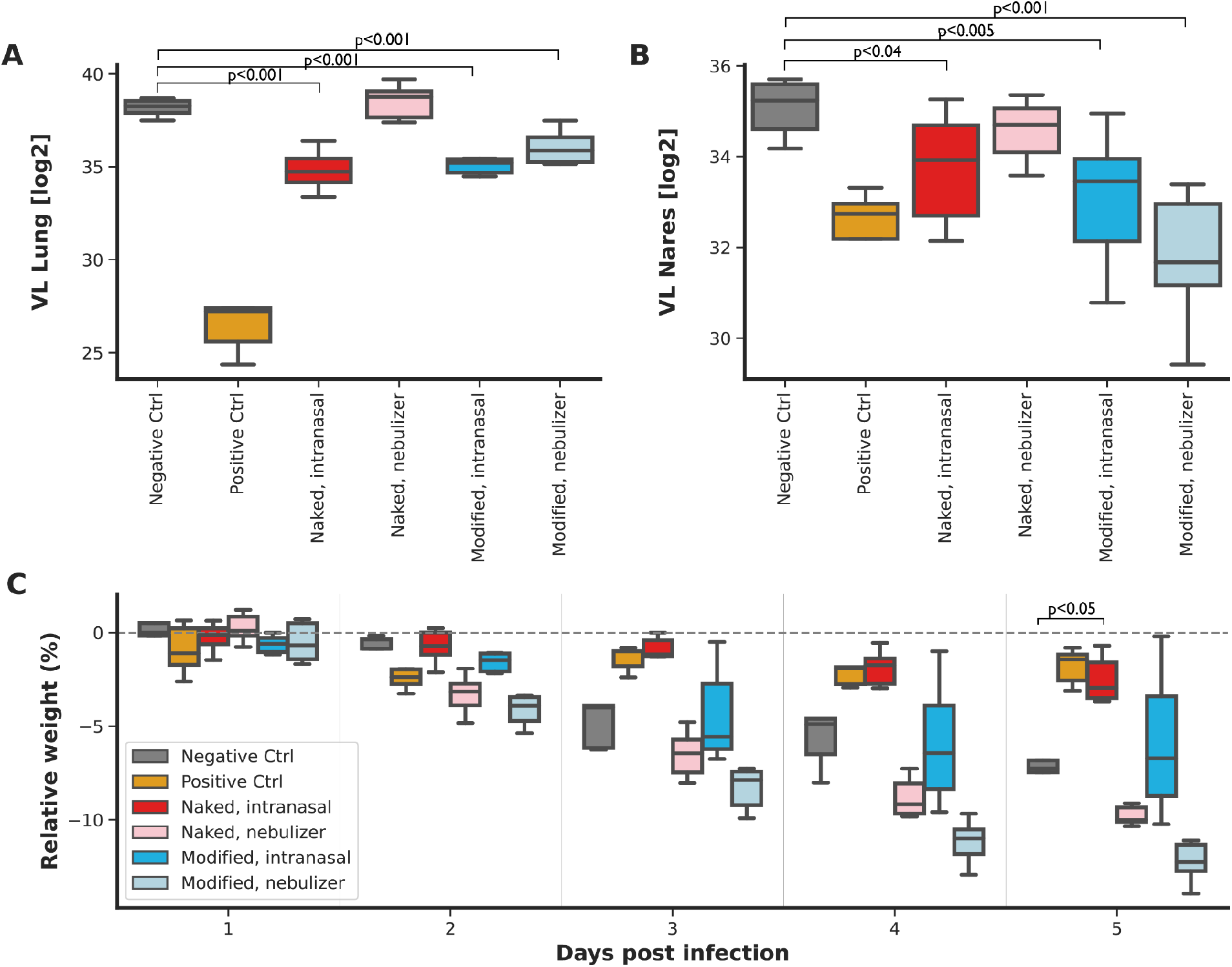
Prophylactic treatments of SARS-CoV-2 infection in Syrian hamsters under various formulations. Syrian hamsters pre-treated with a non-targeting siRNA (negative control; grey), the LY-CoV555 antibody (positive control; orange) or our lead siRNA cocktail in four different formulations, all of which containing our delivery moiety: naked siRNAs intranasally (red), nebulized naked siRNAs (pink), modified siRNA intranasally (blue), nebulized modified siRNA (light blue) **(A-B) Viral load (VL) by qPCR.** All measurements are based on qPCR of the RdRP gene five days post infection from either homogenised **(C) Weight change after infection**. The box plot presents the change in weight by treatment group relative to the infection day. All p-values are adjusted to multiple hypotheses of the four treatment arms and are based on a parametric bootstrapping as described in **Methods**.

**Supplementary Table 1.**
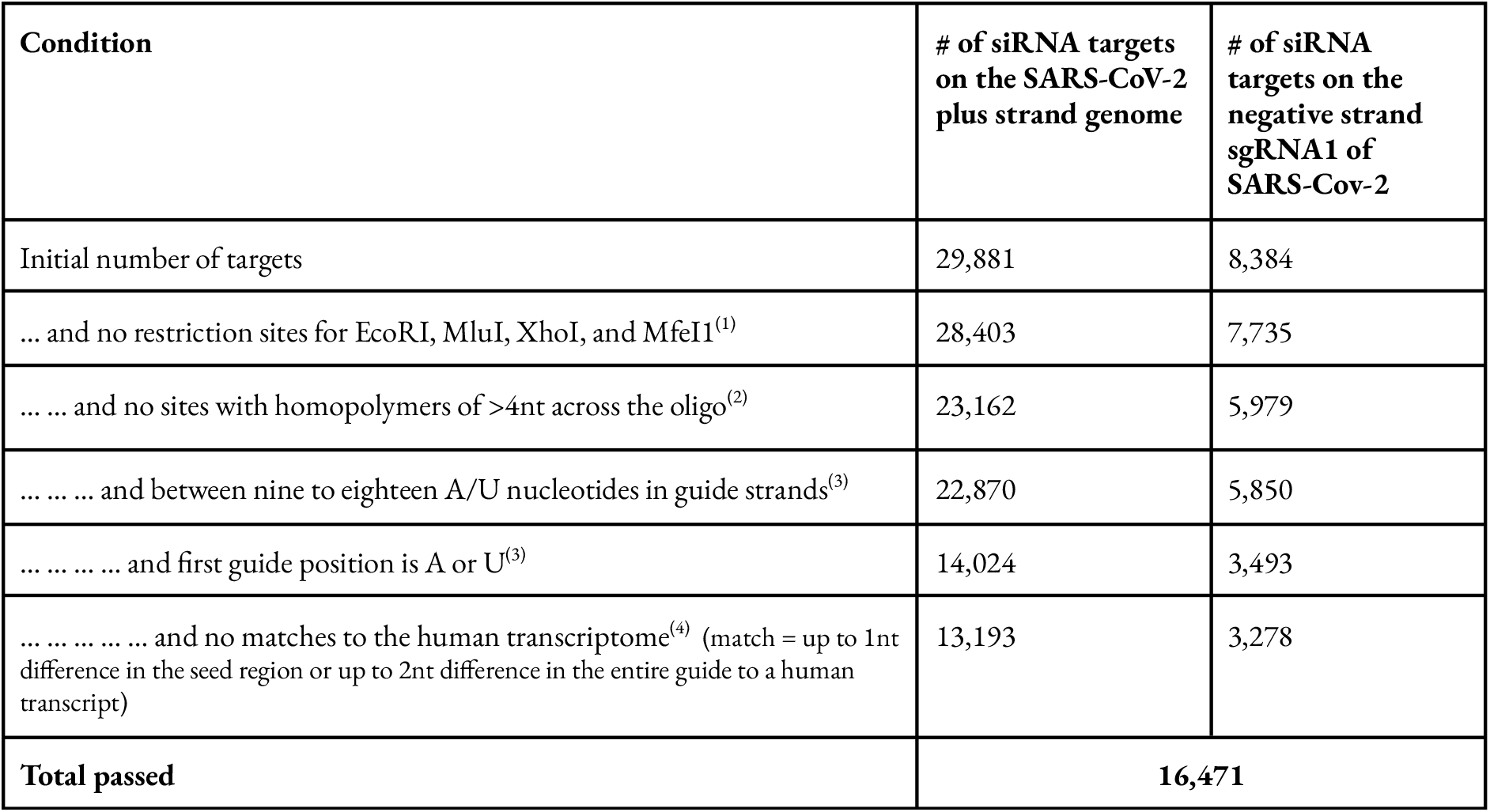
The filtering steps for initial selection of siRNA targets. (1) The restriction enzymes in our cloning strategy. Oligos with these sites will be cut in the middle and therefore are unclonable (2) previous studies^18^ have shown that long homopolymers above 4nt are likely to introduce synthesis, PCR, and sequencing errors. Therefore, we excluded such sites (3) Previous work^13^ has identified these rules of thumb as correlating with potent shRNA response (4) We selected these rules to reduce the likelihood of off-target effects to the human transcriptome.

**Supplementary Table 2.** Replicate-specific IC50 estimates of the siRNA candidates. For each siRNA candidate, we report its IC50 estimate (and its standard error) in replicate 1 (IC50-1), in replicate 2 (IC50-2), and in the analysis that considers both replicates (IC50-both).

| Name | IC50-1 | IC50-1 s.e. | IC50-2 | IC50-2 s.e. | IC50-both | IC50-both s.e. |
| --- | --- | --- | --- | --- | --- | --- |
| **S1** | 41.1 | 5.0 | 35.4 | 3.4 | 38.9 | 3.8 |
| **S2** | 17.3 | 1.1 | 14.4 | 0.8 | 16.0 | 0.9 |
| **S3** | 10.1 | 0.5 | 11.5 | 0.6 | 10.7 | 0.6 |
| **S4** | 17.6 | 1.3 | 15.5 | 0.7 | 16.6 | 0.7 |
| **S5** | 27.9 | 1.9 | 27.9 | 2.0 | 27.9 | 2.0 |
| **S6** | 49.4 | 2.9 | 37.5 | 2.0 | 42.6 | 1.9 |
| **S7** | 254.9 | 52.0 | 264.5 | 44.7 | 259.7 | 45.4 |
| **S8** | 9.4 | 0.4 | 8.0 | 0.3 | 8.6 | 0.3 |
| **S9** | 1291.5 | 79.4 | 1703.1 | 77.9 | 1469.5 | 73.5 |
| **S10** | 21.9 | 1.7 | 17.0 | 1.4 | 19.3 | 1.4 |

## References

1. An open letter by a group of public health experts, clinicians & scientists. Covid-19: An urgent call for global ‘vaccines-plus’ action. BMJ 376, o1 (2022).

2. Krammer, F. SARS-CoV-2 vaccines in development. Nature 586, 516–527 (2020).

3. Dong, Y. et al. A systematic review of SARS-CoV-2 vaccine candidates. Signal Transduct Target Ther 5, 237 (2020).

4. Wang, Z. et al. mRNA vaccine-elicited antibodies to SARS-CoV-2 and circulating variants. Cold Spring Harbor Laboratory 2021.01.15.426911 (2021) doi:10.1101/2021.01.15.426911.

5. Greaney, A. J. et al. Comprehensive mapping of mutations to the SARS-CoV-2 receptor-binding domain that affect recognition by polyclonal human serum antibodies. Cold Spring Harbor Laboratory 2020.12.31.425021 (2021) doi:10.1101/2020.12.31.425021.

6. Baum, A. et al. Antibody cocktail to SARS-CoV-2 spike protein prevents rapid mutational escape seen with individual antibodies. Science 369, 1014–1018 (2020).

7. Muik, A. et al. Neutralization of SARS-CoV-2 lineage B.1.1.7 pseudovirus by BNT162b2 vaccine-elicited human sera. Cold Spring Harbor Laboratory 2021.01.18.426984 (2021) doi:10.1101/2021.01.18.426984.

8. Wu, K. et al. mRNA-1273 vaccine induces neutralizing antibodies against spike mutants from global SARS-CoV-2 variants. Cold Spring Harbor Laboratory 2021.01.25.427948 (2021) doi:10.1101/2021.01.25.427948.

9. Chemaitelly, H. et al. Waning of BNT162b2 Vaccine Protection against SARS-CoV-2 Infection in Qatar. N. Engl. J. Med. 385, e83 (2021).

10. Levine-Tiefenbrun, M. et al. Waning of SARS-CoV-2 booster viral-load reduction effectiveness. bioRxiv (2021) doi:10.1101/2021.12.27.21268424.

11. Ferdinands, J. M. et al. Waning 2-Dose and 3-Dose Effectiveness of mRNA Vaccines Against COVID-19-Associated Emergency Department and Urgent Care Encounters and Hospitalizations Among Adults During Periods of Delta and Omicron Variant Predominance - VISION Network, 10 States, August 2021-January 2022. MMWR Morb. Mortal. Wkly. Rep. 71, 255–263 (2022).

12. Gazit, S. et al. Relative effectiveness of four doses compared to three dose of the BNT162b2 vaccine in Israel. bioRxiv (2022) doi:10.1101/2022.03.24.22272835.

13. Gagne, M. et al. mRNA-1273 or mRNA-Omicron boost in vaccinated macaques elicits similar B cell expansion, neutralizing antibodies and protection against Omicron. Cell (2022) doi:10.1016/j.cell.2022.03.038.

14. Ying, B. et al. Boosting with Omicron-matched or historical mRNA vaccines increases neutralizing antibody responses and protection against B.1.1.529 infection in mice. doi:10.1101/2022.02.07.479419.

15. Sun, J. et al. Association Between Immune Dysfunction and COVID-19 Breakthrough Infection After SARS-CoV-2 Vaccination in the US. JAMA Intern. Med. (2021) doi:10.1001/jamainternmed.2021.7024.

16. Kemp, S. A. et al. SARS-CoV-2 evolution during treatment of chronic infection. Nature 592, 277–282 (2021).

17. Corey, L. et al. SARS-CoV-2 Variants in Patients with Immunosuppression. N. Engl. J. Med. 385, 562–566 (2021).

18. Li, B.-J. et al. Using siRNA in prophylactic and therapeutic regimens against SARS coronavirus in Rhesus macaque. Nat. Med. 11, 944–951 (2005).

19. Tompkins, S. M., Lo, C.-Y., Tumpey, T. M. & Epstein, S. L. Protection against lethal influenza virus challenge by RNA interference in vivo. Proc. Natl. Acad. Sci. U. S. A. 101, 8682–8686 (2004).

20. DeVincenzo, J. et al. A randomized, double-blind, placebo-controlled study of an RNAi-based therapy directed against respiratory syncytial virus. Proc. Natl. Acad. Sci. U. S. A. 107, 8800–8805 (2010).

21. Bitko, V. & Barik, S. Respiratory viral diseases: access to RNA interference therapy. Drug Discov. Today Ther. Strateg. 4, 273–276 (2007).

22. Fellmann, C. et al. Functional identification of optimized RNAi triggers using a massively parallel sensor assay. Mol. Cell 41, 733–746 (2011).

23. Tan, X. et al. Tiling genomes of pathogenic viruses identifies potent antiviral shRNAs and reveals a role for secondary structure in shRNA efficacy. Proc. Natl. Acad. Sci. U. S. A. 109, 869–874 (2012).

24. Banaszynski, L. A., Chen, L.-C., Maynard-Smith, L. A., Ooi, A. G. L. & Wandless, T. J. A rapid, reversible, and tunable method to regulate protein function in living cells using synthetic small molecules. Cell 126, 995–1004 (2006).

25. Love, M. I., Huber, W. & Anders, S. Moderated estimation of fold change and dispersion for RNA-seq data with DESeq2. Genome Biol. 15, 550 (2014).

26. Pelossof, R. et al. Prediction of potent shRNAs with a sequential classification algorithm. Nat. Biotechnol. 35, 350–353 (2017).

27. Tafer, H. et al. The impact of target site accessibility on the design of effective siRNAs. Nat. Biotechnol. 26, 578–583 (2008).

28. Vert, J.-P., Foveau, N., Lajaunie, C. & Vandenbrouck, Y. An accurate and interpretable model for siRNA efficacy prediction. BMC Bioinformatics 7, 520 (2006).

29. Lu, Z. J. & Mathews, D. H. OligoWalk: an online siRNA design tool utilizing hybridization thermodynamics. Nucleic Acids Res. 36, W104–8 (2008).

30. Zheng, B.-J. et al. Prophylactic and therapeutic effects of small interfering RNA targeting SARS-coronavirus. Antivir. Ther. 9, 365–374 (2004).

31. Akerström, S., Mirazimi, A. & Tan, Y.-J. Inhibition of SARS-CoV replication cycle by small interference RNAs silencing specific SARS proteins, 7a/7b, 3a/3b and S. Antiviral Res. 73, 219–227 (2007).

32. He, M.-L. et al. Kinetics and synergistic effects of siRNAs targeting structural and replicase genes of SARS-associated coronavirus. FEBS Lett. 580, 2414–2420 (2006).

33. Zhao, H. et al. SARS-CoV-2 Omicron variant shows less efficient replication and fusion activity when compared with Delta variant in TMPRSS2-expressed cells. Emerg. Microbes Infect. 11, 277–283 (2022).

34. Berkyurek, A. C. et al. A scalable pipeline for SARS-CoV-2 replicon construction based on de-novo synthesis. bioRxiv 2022.02.05.478644 (2022) doi:10.1101/2022.02.05.478644.

35. Huang, H. et al. Profiling of mismatch discrimination in RNAi enabled rational design of allele-specific siRNAs. Nucleic Acids Res. 37, 7560–7569 (2009).

36. Schwarz, D. S. et al. Designing siRNA that distinguish between genes that differ by a single nucleotide. PLoS Genet. 2, e140 (2006).

37. Becker, W. R. et al. High-Throughput Analysis Reveals Rules for Target RNA Binding and Cleavage by AGO2. Mol. Cell 75, 741–755.e11 (2019).

38. Roy, C. et al. Trends of mutation accumulation across global SARS-CoV-2 genomes: Implications for the evolution of the novel coronavirus. Genomics 112, 5331–5342 (2020).

39. Chen, P. et al. SARS-CoV-2 Neutralizing Antibody LY-CoV555 in Outpatients with Covid-19. N. Engl. J. Med. 384, 229–237 (2021).

40. Rosenke, K. et al. Defining the Syrian hamster as a highly susceptible preclinical model for SARS-CoV-2 infection. Emerg. Microbes Infect. 9, 2673–2684 (2020).

41. Chan, J. F.-W. et al. Simulation of the Clinical and Pathological Manifestations of Coronavirus Disease 2019 (COVID-19) in a Golden Syrian Hamster Model: Implications for Disease Pathogenesis and Transmissibility. Clin. Infect. Dis. 71, 2428–2446 (2020).

42. Yuan, L. et al. Gender associates with both susceptibility to infection and pathogenesis of SARS-CoV-2 in Syrian hamster. Signal Transduct Target Ther 6, 136 (2021).

